# Automated Stimulation and Long-Term Remote Monitoring of Multi-Plex Microfluidic Organoid Chips

**DOI:** 10.1101/2023.10.02.560607

**Authors:** David M Sachs, Kevin D Costa

## Abstract

Developing a novel microfluidic organoid system required many experiments and iterations due to lack of knowledge about the relevant developmental biology. Collecting data on the developing organoids quickly escalated into a bottleneck as high throughput and long term culture resulted in a rapidly increasing number of specimens being observed. Commercially available automated microscope systems exist, but were either too expensive or not appropriate, and could not be modified. To satisfy the increasing need for automated data collection, a custom robotic system was developed to collect data from within a standard incubator. An X-Y belt driven gantry was designed with an architecture chosen to balance high accuracy, low cost, speed, and range of motion. Focus control was implemented with dual miniature leadscrews. A linear sliding mechanism was used to switch between two microscope objectives. 3D printed chip attachments were designed to implement illuminators for bright field imaging, and electrodes for stimulating the cardiac organoids. A fluorescent filter block was designed using a 3D printed piece to hold optical components, and a multi-band filter set that allowed for three color fluorescence without moving parts. A pulley driven tilting stage gravitationally biased the organoids during development. In order to ensure accurate image collection despite the inevitable position shifting of the chips, an image processing pipeline was developed for locating organoids using geometrical microfluidic chip features. The resulting robotic system automated imaging data collection on organoids and electrical and mechanical stimulation, in addition to being modifiable for future projects.

## Introduction

High throughput monitoring of developing organoid systems benefits from automation, especially when differentiation protocols are uncertain and still being optimized. Live imaging within an incubator allows for less invasive monitoring, maximizing incubation stability and minimizing the handling of sensitive organoid systems. In addition, for cardiovascular organoids, electrical pacing of cardiomyocytes is dependent on temperature, and benefits from being performed in an incubated environment.^1^ Motorized microscopes that can function within incubators are commercially available^2-6^ or described in published literature,^7-33^ but may not have the necessary price point or features for widespread use in an academic research setting. While expensive research equipment is often acquired by institutions and shared among labs, equipment for time lapse imaging within an incubator over months-long time scales is not something that can easily be shared.

While some popular microscopes contain motorized stages and objective turrets that allow for automation, such as the Keyence BZ-X800^34^ and the EVOS M700,^35^ these microscopes are used outside incubators, and rely on users to carry plates to them and manually locate desired fields of view. Other commercial products enable live cell imaging within incubators, such as the Holomonitor^4^ and Etaluma LS720,^3^ but these are low throughput systems. The Olympus VivaView^5^ implements automated microscopy through the bottom of a modified multi-gas incubator; however, this system is limited to the imaging of six 35 mm dishes, and the product line has been discontinued. The CellCyte is a recently released product that is close to the system required of this project; however, in addition to costing at least $50k^2^ it isn’t modifiable and lacks some necessary features, such as high frame rate video for imaging beating cardiac organoids and flow, as is required for this application.

More sophisticated live cell imaging systems may incorporate robotic plate handlers to move between incubators and microscopes, such as the Biotek Biospa and Cytation.^6^ These systems enable high throughput time lapse imaging, but a retail cost of about $150k is outside the budget of a typical academic biology lab, and they involve considerable manipulation of samples as they leverage plate handling to remove plates from a custom incubator and pass them to a microscope.

Some labs with technical expertise have developed and published their own microscope systems.^7-33^ Custom systems may not have the sophistication or image quality of a commercial system, but can be much cheaper and are modifiable for specific applications. While optical components remain expensive to both commercial and academic microscope developers, electronic and mechanical components can be obtained cheaply due to the vast economic scale of consumer electronics and home appliances, and the recent boom of open source 3D printers provides a wealth of options for gantry design and motor control.

Some low cost DIY microscope systems are designed for portability for point-of-care solutions to specific problems, such as cytometry,^7-9^ imaging for diagnostics,^10-15^ parasite classification,^11, 16, 17^ agricultural field work,^18^ electrophysiology,^19-21^ or education.^22-27^ Such systems provide useful guidance for designing microscopes, but make image quality sacrifices in order to meet portability and cost requirements. While complete motorized microscopes might be outside the budget of a small lab, the optical components themselves, such as complex multiband fluorescence filters, may be affordable. Gantry accuracy and optical component alignment might be compromised in the effort to implement an affordable DIY system, but it shouldn’t be necessary to compromise the quality of the optical components themselves.

Other microscope projects are intended for live cell imaging within incubators,^28-33^ but don’t have the mechanical range or design features required by this project. Some DIY microscopes make use of existing 3D printer gantries in order to easily implement motor control without designing a new, custom gantry;^36, 37^ however, such projects have limited fields of view due to the small print beds used by 3D printers, and the overall inefficiency of integrating a microscope into a gantry that was designed for a different application.

For monitoring microfluidic cardiovascular organoids, the design features required were: 1) bright field and fluorescence capabilities for monitoring tissue morphology and flow tracers respectively, 2) at least 60 frames per second video for capturing cardiomyocyte contractions and flow patterns, 3) magnification such that high quality imaging of the 800 μm diameter central organoid wells can be acquired, 4) a second higher magnification for capturing fluorescence flow tracers in channels with less background fluorescence, 5) a range of motion that is a big as possible in order to fit as many chips on the stage as possible, 6) image processing for the robot to find the correct fields of view even if the chips have shifted due to use by biologists, 6) the ability to program the robot in order to extend its capabilities, and 7) a tilting stage, as the cardiovascular organoids are cultured at a 30 degree angle. The last requirement is clearly problematic for the typical bright field imaging system, as the bright field illuminator would make it impossible to move the stage.

Due to the lack of a commercial or published robotic microscope with all the necessary features required by this project, and the advantages of being able to modify the system as needed, a custom system was developed.

## Methods

### Machine Design and Construction

Machines were modelled in Fusion 360 (Autodesk). Off-the-shelf parts were acquired from McMaster-Carr or Misumi, with custom parts 3D printed in ABS on an Ultimaker S3 3D printer or laser cut in acrylic on an Epilog Helix. Low level control of sensors, actuators, and LEDs was performed on Arduino Neros (Digi-Key), with separate Arduinos controlling the motors and electrical pacing. Gantry and tilter NEMA-17 stepper motors were controlled by ST L6470 motor drivers on X-NUCLEO-IHM02A1 boards (Digi-key), with the boards configured to be stacked in a daisy chained manner.

### Software Design and Architecture

Arduino firmware was used to perform low level interfacing to actuators, sensors, and LEDs, while accepting serial commands and outputting a stream of motor location and sensor data. Arduinos were connected to a desktop machine running custom software on Windows 10 that integrated control of the motors, electrical pacing, and microscope imaging and lighting into one C++ program. OpenCV was used for real time image processing in order to track features and automate time lapse protocols via a custom G-Code like scripting language, while text commands were provided for real time driving of the system.

## Results

### Gantry

For mechanized microscopes for live cell imaging, the ability to return to the same field of view repeatedly is a requirement. This problem is typically solved by making the gantry as accurate as possible. Producing a low-cost gantry with 3D printed parts capable of both a wide range of motion, high speed, and high repeatability would be a daunting task. However, because the long term tracking of differentiating and developing organoids also requires that users be able to remove and reposition microfluidic chips, gantry repeatability is not a viable solution to the problem of imaging repeatability, as the position and orientation of the chips themselves will not be precisely repeatable; locating organoid positions would be a task for machine vision software aided by mechanical gantry accuracy.

The cost of the gantry and microscope are intimately tied to the application. In our case, the application is the long-term imaging of organoids within microfluidic chips that are accessible to the user and might not be repositioned accurately. The user accessibility requirement eliminates simple mechanical repeatability as the sole solution to organoid tracking, but the microfluidic structures simplify the task of image tracking with computer vision algorithms. As computer vision is an attractive solution to time-lapse imaging for this application, the burden of mechanical repeatability is lifted from the gantry and the microscope. If mechanical repeatability were necessary, such a robotic system would require high-cost bearings, steel structural components, and rigid optical interfaces. The shift from reliance on precision hardware to reliance on smart software enables the use of low-cost bearings and less rigid 3D printed structural components and optical fixtures that may shift positions over time.

The X-Y gantry structure carrying the robotic microscope was designed to maximize the range in X and Y in order to maximize the number of organoids that could be observed. Some candidate gantries are shown in **Figure 1A-C** along with the final implemented design in **Figure 1D**. While the single plane H architecture (**Figure 1A**) allows for a wide X-Y range and a low profile, it suffers from racking inaccuracies at high accelerations due the coupling between X and Y motion,^38^ and such high accelerations might be advantageous in an automated time lapse microscope intended to minimize imaging time. The low profile is less useful in this case because the height is constrained by the microscope.

**Figure 1.**
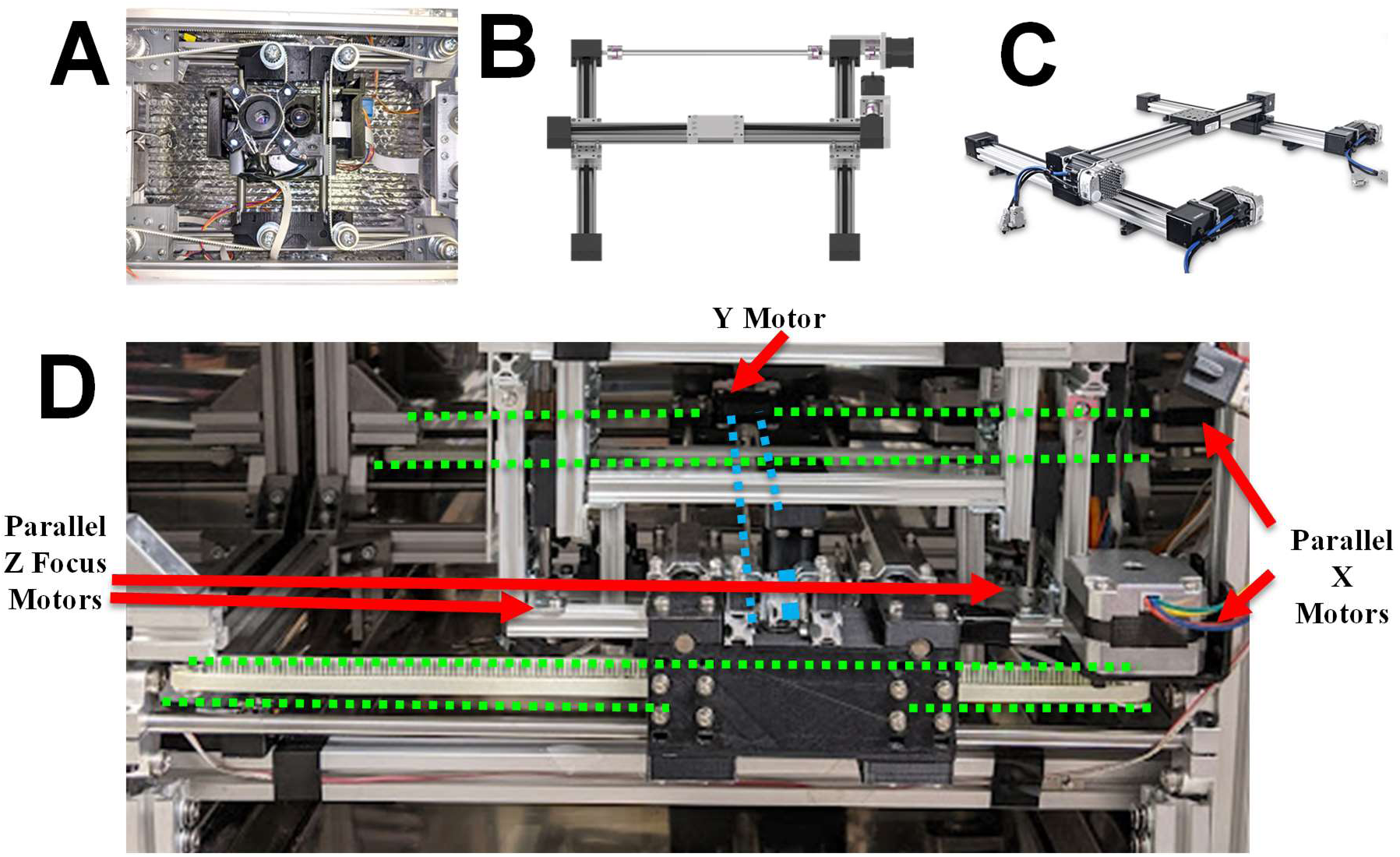
Gantry Candidates and Final Design. **A)** Single plane parallel H-bot gantry **B)** Dual plane serial H gantry with axle coupling X belts, adapted from Igus. **C)** Dual plane H gantry with X driven by two motors, adapted from Newmark. **D)** Parallel X motors move the gantry along the X-axis via two belt mechanisms (green dashed lines) while a serial Y motor moves the gantry along the Y-axis via a decoupled belt mechanism (blue dashed line), and two parallel Z motors focus the microscope via parallel miniature leadscrews.

Two other H like gantries are shown in **Figure 1**: an implementation shown by Igus^39^ (**Figure 1B**) uses two planes and two motors, and couples the two X mechanisms by an axle, whereas an implementation by Newmark^40^ (**Figure 1C**) eliminates the axle by using two separate X motors. This three motor H architecture has a disadvantage of requiring three motors to control two axes, and is therefore overconstrained and risks binding if the two X motors become unsynchronized. However, if the two motors remain synchronized, this three motor architecture is simple, precise, and stable,^41^ and reduces the burden on the frame to be rigid, as the frame is not transmitting a force from one side to the other via the axle. In addition, the removal of the axle opens up valuable space that increases the range of movement of the microscope. The implementation chosen was this three motor gantry architecture (**Figure 1D**).

Similarly, the Z focus mechanism was designed to use two motors driving two flanking miniature leadscrews (Misumi MSSR601-100), allowing the microscope to have a large set of bulky features without a stiff, steel frame (**Figure 1D** and **Figure 2**).

**Figure 2.**
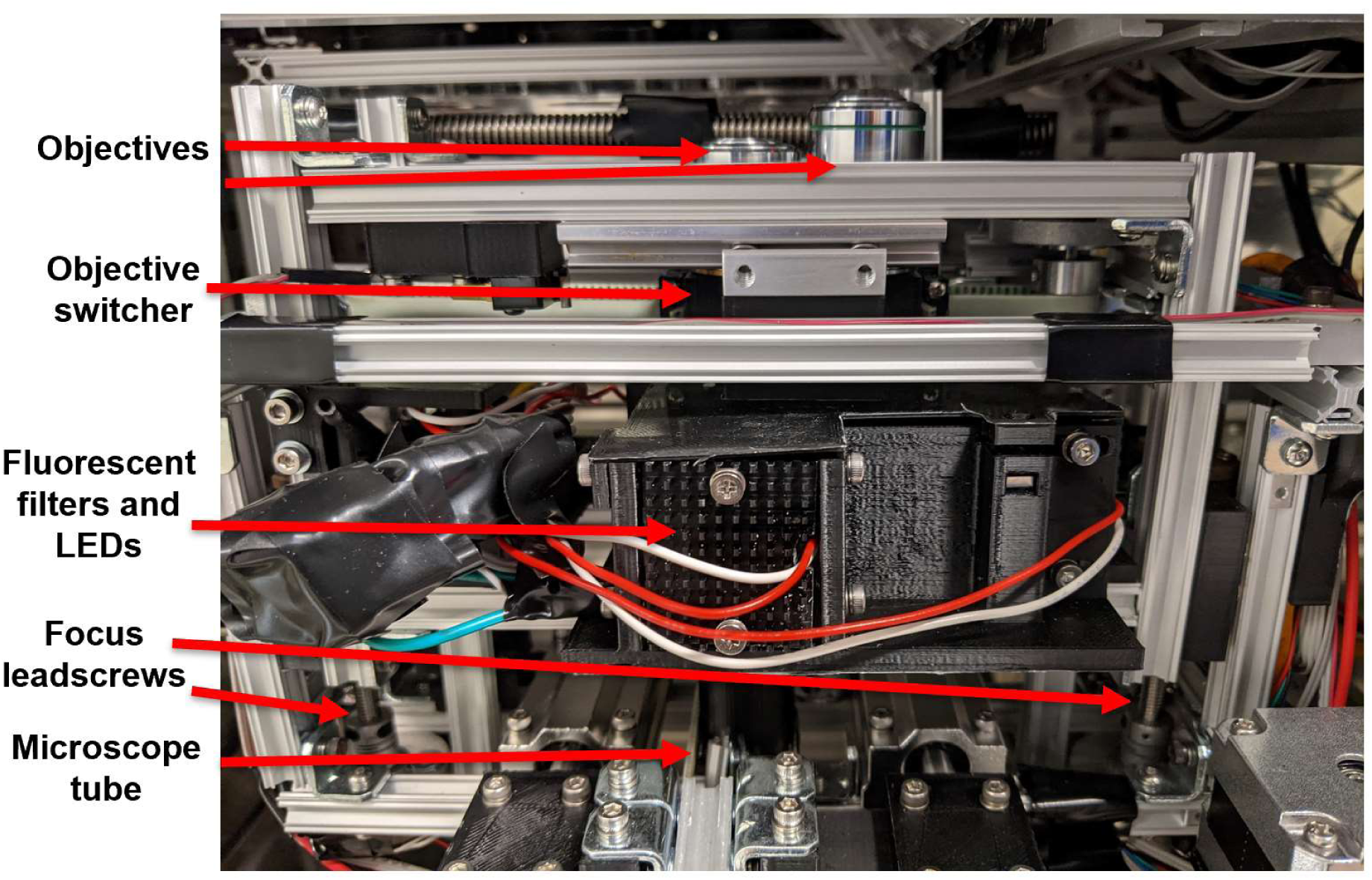
Microscope. Aluminum frame holding microscope optics and mechanisms, including dual objective switcher, fluorescent filter block and LEDs, focusing motors and leadscrews, and microscope tube with tube lens and USB3 camera.

### Microscope

The microscope itself used a belt mechanism and sliding lens mount to switch between two objectives, which were typically 10X and a 20X objectives (**Figure 2**). The field of view through the 10X objective was 1620 μm by 900 μm when the camera was in a 1280×720 pixel mode, corresponding to about 1.3 μm per pixel, and was chosen to be slightly larger than the 800 μm organoid wells in the microfluidic chips, maximizing the resolution while enabling a single frame to capture the entire organoid along with a small field of view of the connecting channels. The 20X objective was chosen to achieve a closer view of the channels, and for decreasing the background fluorescence when fluorescent imaging is used.

The microscope used a 50 mm focal length tube lens (Thorlabs AC254-050-A) transmitting images to a USB 3 camera capable of 60 fps monochrome 1280×720 pixel frames with a global shutter (Leopard Imaging LI-USB30-M021M) (**Figure 2**). The camera was also capable of 30 fps capture with an image size increased to 1280×960 pixels, or 90 fps capture with an image size reduced to 800×460 pixels. The global shutter was critical for eliminating the rolling shutter artifacts that would be generated by more widely available CMOS cameras when imaging beating cardiac organoids.^42^ The 50 mm tube lens was adapted from the OpenFlexure microscope,^43^ and allows for a compact microscope tube design compared with the more typical 160 mm – 200 mm microscope tube lengths, but at the cost of higher chromatic aberration, producing slightly blurrier images if illuminated with white light. This problem was remedied by using single wavelength 530 nm LEDs (Mouser, 416-LST101F06GRN1) to illuminate the chips during bright field images, with **Figure 3** showing comparison images between the green LED and a white LED (Mouser 749-R20WHT-F-0160). To compare the two, cells were imaged in a chip that undergone a differentiation that had led to monolayers in some regions of the central organoid wells. The objective switcher was set to 20X, and a region of interested was focused on using the green LED (**Figure 3A**). The LED was then swapped out with the white LED, and an attempt was made to get a comparatively focused image (**Figure 3B**). The image from the green LED is noticeably sharper when attempting to focus on individual cells. Note that this imaging was executed through a triple-band bandpass filter, as described below, so “white” actually means three specific frequencies: 446 nm, 532 nm, and 646 nm. Using a single color LED additionally reduced phototoxicity, as the bandpass filter blocked most of the power from the white LED, requiring the cells to be exposed to an unnecessarily high brightness in order for a subset of the light to reach the camera. With the 530 nm LED, chosen to be compatible with the bandpass filter, most of the light exposing the cells reached the camera.

**Figure 3.**
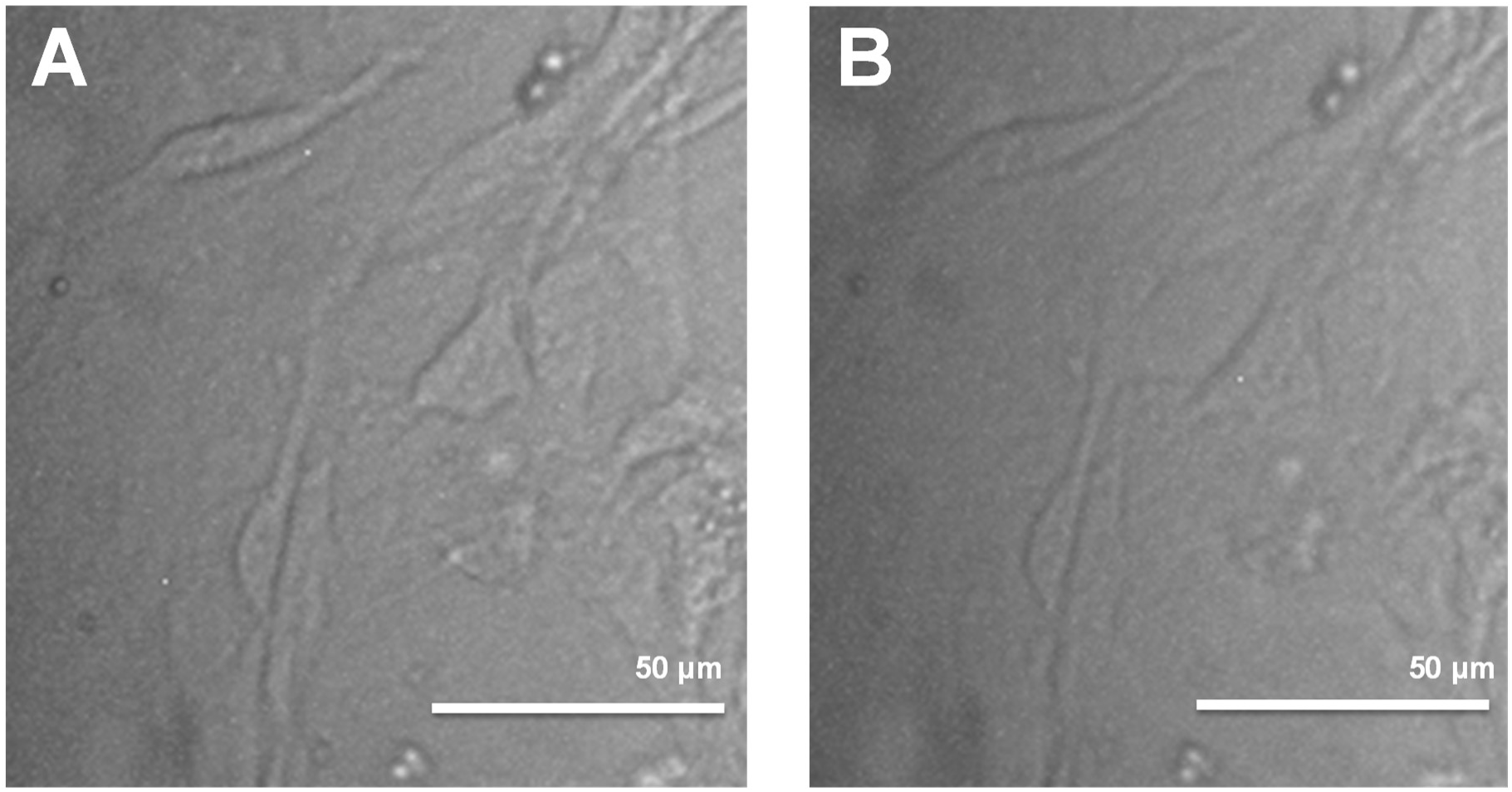
Single Color Bright Field Imaging. **A)** Bright field image of a monolayer of cells with the 20X objective and a green LED. **B)** The same, but with a white LED.

To enable multi-color fluorescent imaging without the need for additional motors to switch filters, a Pinkel filter configuration^44^ was used that was adapted from the Etaluma LS720.^3^ For the emission filter, a 446/532/646 nm triple-band bandpass filter was used (Semrock FF01-446/532/646-25), and for the dichroic filter a 405/488/594 nm triple-edge dichroic beamsplitter was used (Semrock Di01-R405/488/594-25). Separate excitation filters for the LEDs were used, with a 390 nm filter (FF01-390/40-25) for the 390 nm LED (LED Engin LZC-C0UB00-00U4), a 482 nm filter (Semrock FF02-482/18-25) for the 485 nm LED (New Energy XPEBBL-L1-0000-00301-SB01), and a 591 nm filter (Semrock FF01-591/6-25) for the 590 nm LED (LED Engin LZ4-40A108-0000). The LEDs were focused on the back plane of the objective with Fresnel lenses (Thorlabs FRP125), chosen because of their compactness relative to glass lenses, and their light was combined by joining the 390 nm light with the 485 nm light using a 458nm dichroic (Semrock FF458-Di02-25×36), and then joining that with the 591 nm light using a 552 nm dichroic (Semrock FF552-Di02-25×36). The result of this design is that three-color fluorescent imaging is possible without moving parts, with the fluorescent color chosen by illuminating with one of the three LEDs (**Figure 4A**). Additionally, the 390 nm LED could be refocused onto the image plane in order to selectively photoinitiate hydrogels or cleave photoactivable reagents while the chip is being imaged.

**Figure 4.**
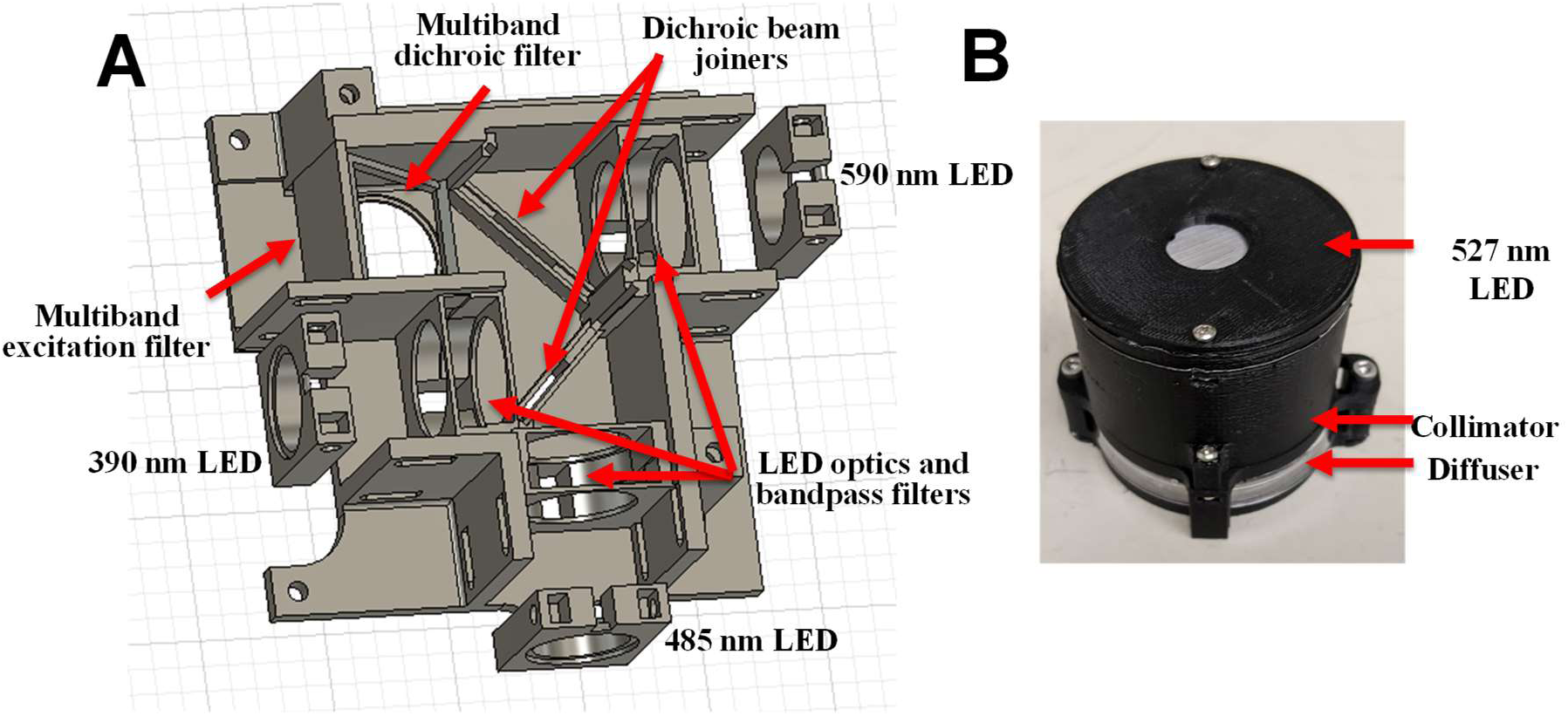
Fluorescent/Bright Field Microscope Design. **A)** Single piece of 3D printed black ABS designed to hold the indicated optical components; LEDs were mounted to separate adjustable pieces. **B)** 527 nm bright field illuminator.

Bright field illuminators illuminate with one wavelength, with 530 nm LEDs chosen as it is compatible with the 532 nm line in the emission filter. The brightfield illuminators were designed to be compact and low cost, as one illuminator per chip would be needed, with up to 12 chips per machine. The illuminators additionally contain Fresnel lenses (Thorlabs FRP232), and diffusers (Thorlabs DG20-1500) to collimate the light (**Figure 4B**). The microscope frame had to be adjustable to allow for aligning and focusing of optical components, while maintaining positions and orientations of optical components over time despite gantry accelerations. The frame was assembled from 10 mm X 10 mm miniature aluminum extrusion support beams (McMaster-Carr 1959N1), with lock washers on the connecting screws, maximizing flexibility of component placement while minimizing position shift over time (**Figure 2**). The complex optical component adapters were 3D printed from black ABS to minimize internal light scattering. While it wasn’t practical to allow every optical filter to be adjustable, the high power fluorescence LEDs were mounted to 3D printed pieces that could be adjusted relative to the rest of the filters, allowing for focusing on the back objective plane to minimize the spatial variation of the illumination (**Figure 4A**).

Example images from the custom microscope are shown in **Figure 5**. These include a bright field image of compacting and differentiating mesoderm (**Figure 5A**) with a 10X objective, and three fluorescence images taken using a 20X objective: green fluorescent 1 μm beads used as flow tracers and imaged at 60 fps (**Figure 5B**), red fluorescent 200 nm beads used to image hydrogel structure (**Figure 5C**), and a channel containing differentiated cells stained for endoglin (green) and DAPI (blue) (**Figure 5D**).

**Figure 0.5.**
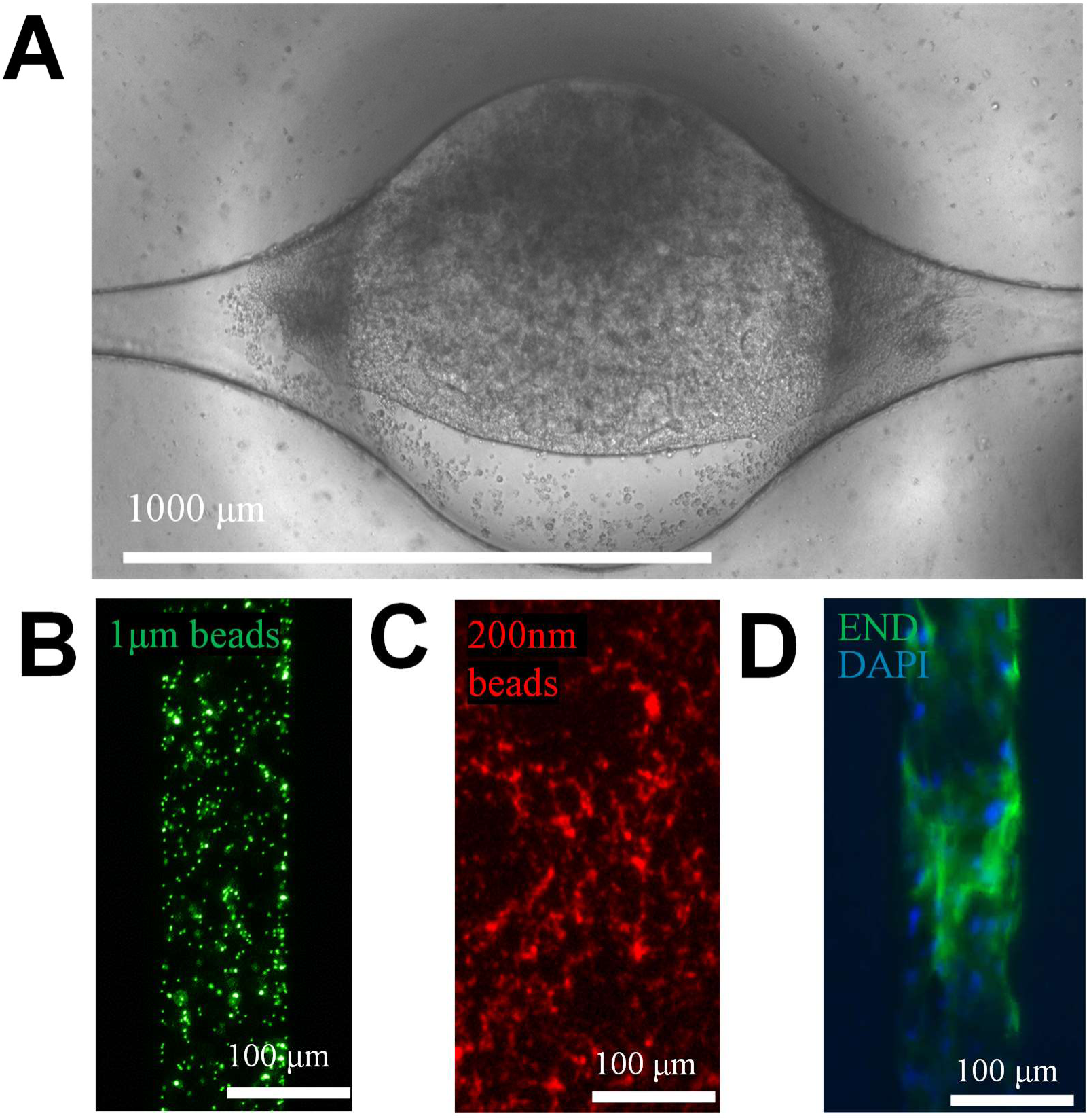
Microscope Image Examples. **A)** Bright field image of differentiating and compacting mesoderm. **B)** Green fluorescent 1 μm beads. **C)** Red fluorescent 200 nm beads. Figure 6**. Dual Tiling Microscope Stages.** Compound pulley mechanism lifted and lowered one end of the microscope stag**e** as accelerometers provided feedback. **D)** Cells in a microfluidic channel stained for endoglin (green) and DAPI (blue).

### Tilter for Perfusion, Cell Biasing, and Flow Impulses

The microscope stage that would hold the microfluidic chips had to be structurally stable while still being thin enough in the vicinity of the samples to allow for objectives to come as close as possible to the target. Long working distance objectives were chosen to maximize the potential thickness of the microscope stage: a 10X objective with a 17.5 mm working distance (Motic AE2000 MET - LM PLAN 10X) and a 20X objective with an 8.1 mm working distance (Motic AE2000 MET - LM PLAN 20X). Future applications may use a higher magnification objective, such as the Motic 50X Objective with an 8.4 mm working distance (Motic AE2000 MET - LM PLAN 50X).

The microscope stage was constructed from a bottom piece of 1/16” thick acrylic chemically welded with SciGrip acrylic cement (McMaster-Carr 7517A2) to a top piece of 1/4” thick acrylic, both cut on an Epilog Helix laser cutter. The thin piece supports the microfluidic chips and minimizes the required working distance, while the thicker piece provides structural support. Shorter working distance objectives would necessitate a thinner stage machined out of steel, and a redesign of the chip, which currently has a thin PDMS layer inside a polystyrene dish positioned between the organoid and the microscope.

In addition to the microscopy requirements for the stage, the stage was designed to tilt in order to provide rocking perfusion to perfusable microfluidic chips, cell biasing in order to sediment cells towards one side of the chip with a fixed 30 degree angle, or small transient flows in order to visualize a flow from a specific pressure head. A tilting mechanism would typically be symmetrical with one side moving upwards while the other side moves downwards, but our design was complicated by the existence of a microscope directly below the stage.

Since the tilter must tilt without any part moving below the focal plane, and then return to a flat position at the focal plane, an asymmetric tilting mechanism was designed, with one end of the stage rotating on a hinge, while the other end is lifted or lowered by a motor-controlled compound pulley mechanism. The tilting stage has an accelerometer providing feedback for both microscope stage angle and vibration. As the desired tilting angle or speed may vary depending on the experiment, two microscope stages were installed with independent tilt mechanisms and control, each holding 6 microfluidic chips mounted in their illuminators (**Figure 6**).

**Figure 6.**
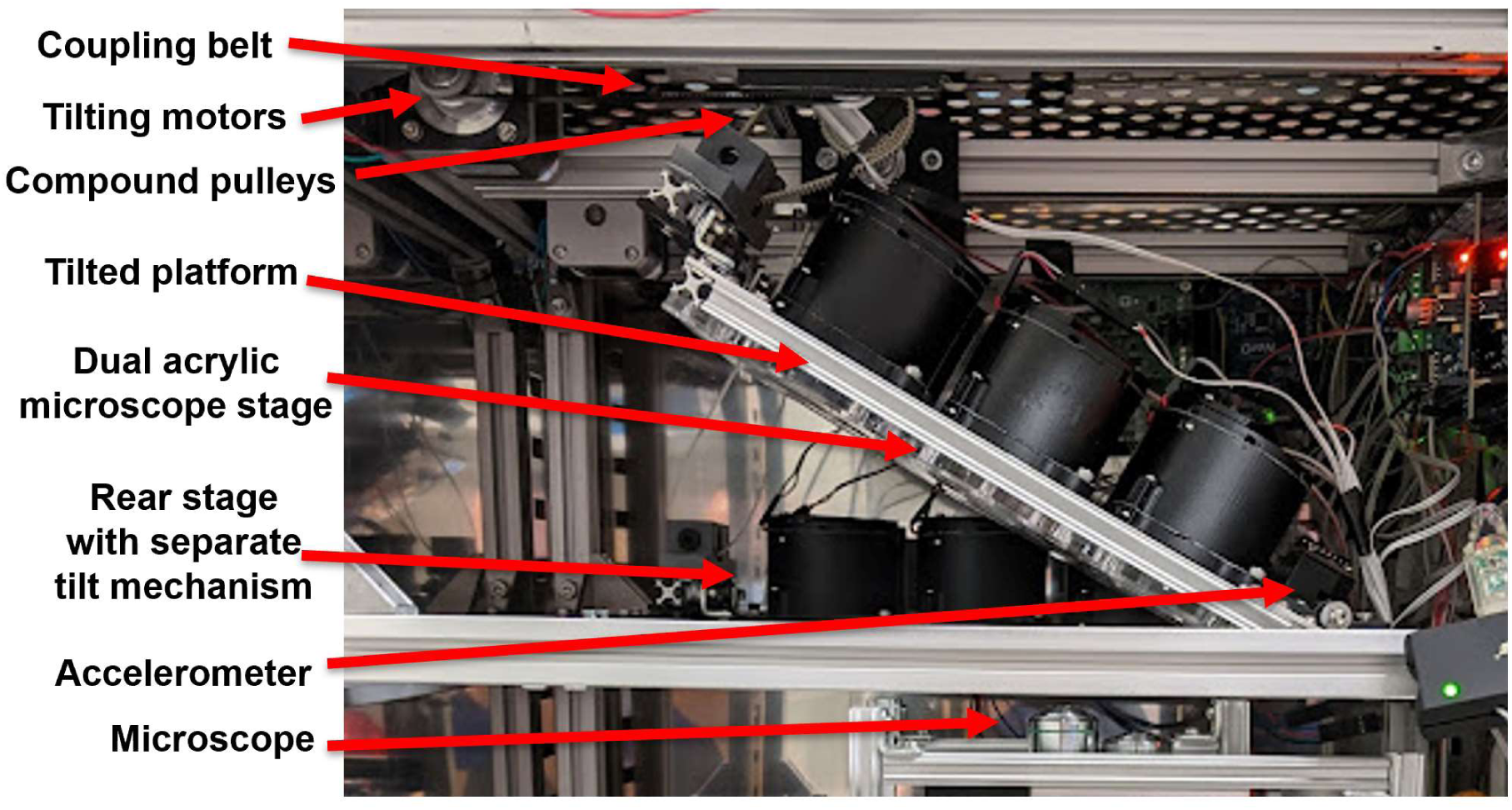
Dual Tiling Microscope Stages. Compound pulley mechanism lifted and lowered one end of the microscope stag**e** as accelerometers provided feedback.

### Electrical Pacing System for Cardiovascular Organoids

Electrical pacing by field stimulation is commonly used to control beat rate when characterizing the contractile function of cardiomyocytes and engineered cardiac tissues. The process involves applying periodic electrical pulses at a range of frequencies and measuring the ability of the cardiac tissue to contract at that frequency. Chronic pacing has also been used to accelerate the phenotypic maturation of engineered cardiac tissue.^45^

Electrodes used for pacing of cardiomyocytes in vitro must be electrochemically inert, and are typically made of graphite,^46, 47^ a low cost, conductive, and inert material that can be purchased in a variety of geometries. By contrast, implantable pacemakers have historically used platinum or platinum-iridium leads for reliable and biocompatible long-term pacing of the heart.^48^ Some microfluidic cardiac models also use platinum wire electrodes.^49^ While graphite electrodes are very effective, they can sometimes shed graphite particles that, though inconsequential to macroscopic engineered tissue strip cultures, could interfere with microfluidic channels. For the cardiac organoid microfluidic chip, 200 μm diameter platinum wire was chosen as the electrode material.

A 3D printed attachment was designed to interface the platinum wire electrodes to the microfluidic chip (**Figure 7A-B**). The attachment was assembled from three thin 3D printed pieces that screwed together to hold both electrodes and wiring. The attachment clamped to the microfluidic chip and sat on top of it, leaving 1 mm of clearance between it and the chip to avoid touching the media in the reservoirs. The low profile of the design allowed it to fit within the 60 mm dish that held the chip, and spaces in the attachment allowed chip media to be exchanged without removing the electrodes.

**Figure 7.**
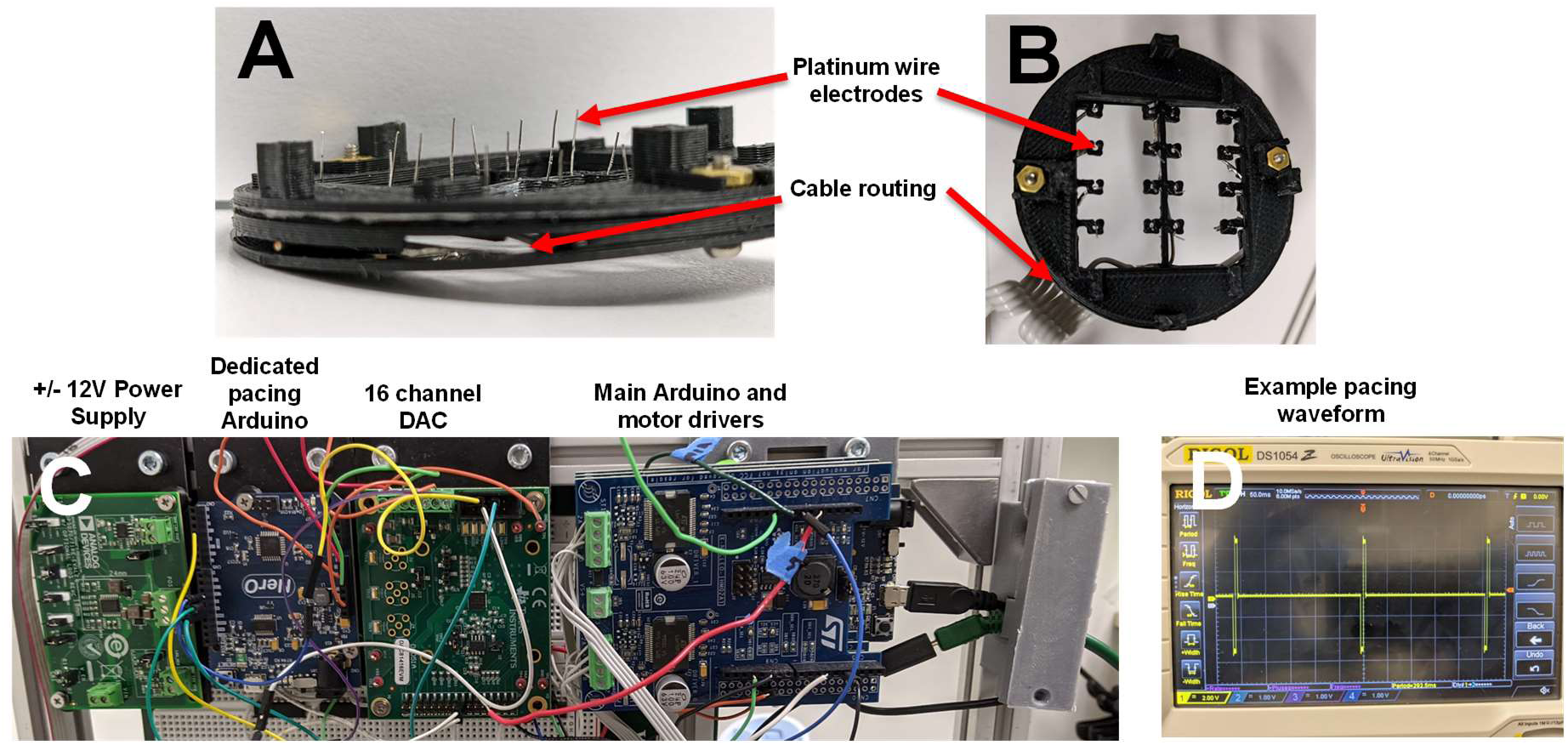
Pacing System. **A)** Platinum wire electrodes were embedded in a 3D printed fixture (upside-down). **B)** Top view of the fixture as it would sit on top of a chip, with electrodes in the reservoirs. **C)** Electronic circuitry, including positive and negative power supplies for biphasic pulses, a dedicated pacing Arduino, a 16 channel DAC, and the main Arduino and motor drivers.

Pacing systems for engineered cardiac tissue generate electrical field gradients of 0.48 V/mm across the cardiomyocytes, with frequencies usually ranging from 0.25 Hz to 2 Hz, but occasionally using higher frequencies.^45, 50^ Pulse widths should be as short as possible to minimize electrochemical modification of the electrodes, and biphasic pulses are similarly used to avoid an average DC current that could result in electrode electrolysis.^51^ The required voltage for the microfluidic organoid chip depended on the distance travelled between the electrodes, which was 20.65 mm as measured from the tops of the reservoirs, resulting in an estimated source voltage of 9.9 V. The current required to drive this voltage was given by the resistance of the microfluidic structures when filled with cell culture media, which has an approximate conductivity of 50 Ω·cm.^52^ The predicted resistance of 18 kΩ was primarily the resistance of the shunt, which was in parallel with the higher resistance of the channel. The current required to drive this system was therefore around 0.5 mA.

The electronics for the pacing system were designed to be as flexible as possible, allowing a variety of programmable waveforms to simultaneously stimulate multiple chips, supporting the use of both chronic and periodic pacing. The pacing system was driven by a dedicated Arduino to ensure that the timing of the waveform wouldn’t be compromised by multitasking of the microcontroller. The Arduino sent SPI commands to a set of 16 channel DACs (DAC81416EVM) capable of +/- 20 V outputs with buffer outputs that could drive 25 mA. The DAC was powered by a positive/negative switching power supply that generated positive and negative rails of up to +/- 39 V (ADP5071RE). The rails were additionally filtered to produce low noise +/-12 V rails generated by low-dropout regulators (LDOs) (ADP5071RE-EVALZ) (**Figure 7C**). The Arduino firmware could generate 16 simultaneous waveforms, which were chosen to be monophasic or biphasic pulses of programmable voltage, frequency, and pulse width (**Figure 7D**).

### Image Processing for Automated Time Lapse

Automating time lapse imaging of the microfluidic organoids required a system that could repeatably return to specific fields of view, focal planes, and illuminations. As the organoid had a diameter of 800 μm, and would ideally fill the field of view in order to maximize the resolution of biological structures, the gantry mechanism needed to be repeatable to within 100 μm. However, this time lapse system was additionally challenged by lack of repeatability in other aspects. The microfluidic chips were removed and replaced daily for manual media changes, with variation in placement position and angle after each access. Old microfluidic chips were regularly swapped out with new chips that had some variation in the X-Y placement of the microfluidic structures and focal plane of the organoid tissue within the chip. Standard autofocus algorithms could pick an undesirable focal plane, as the microfluidic structures had 3D aspects that could be in focus at a variety of planes, and the organoids themselves were also 3D. To handle the automation requirements for this particular time lapse system, a custom image processing pipeline was developed that allowed the machine to locate and image desired microfluidic structures and was robust against variations in chip position and orientation.

The first step in the pipeline was to locate the organoid well. This was an 800 μm circular well, so a circle Hough transform was used to locate it (**Figure 8A-C**). The Hough transform could occasionally be triggered by the curved edges connecting the channels to the wells, combined with noise at the bottom focal plane of the microfluidic structures. The well itself was 2 mm deep, and would generate visible circles at any focal plane within that range, so these false positives could be eliminated by raising the focal plane up by 500 μm, ensuring that only the 2 mm through-holes were in focus (**Figure 8B**). The edge detection algorithm that ran prior to the Hough transform was very sensitive to lighting, so the algorithm began by sweeping through a range of lighting brightness settings and choosing the image with the best circle Hough transform hit. Typically, this occurred when the lighting was bright enough to saturate the noise in other parts of the image, but still allowed the circle to be visible.

**Figure 8.**
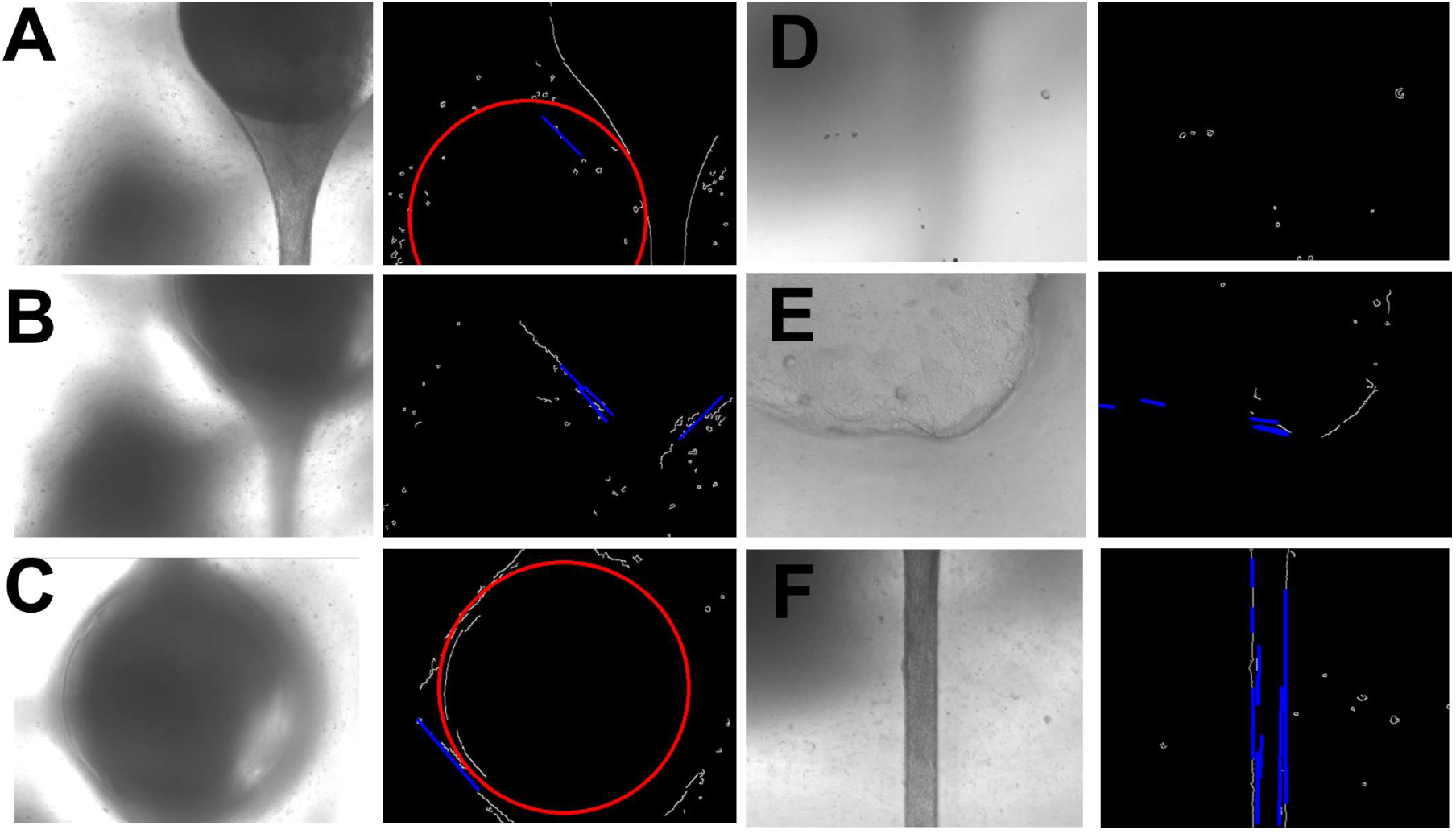
Image Processing for Time Lapse Automation. Bright field images (left) were paired with the OpenCV image analyses (right) showing edges (white), circle Hough transform hits (red) and line Hough transform hits (blue). **A)** A false positive circle hit resulting from a curved edge combined with noise. **B)** The false positive was eliminated by choosing a higher focal plane where the feature no longer exists. **C)** The higher focal plane results in a correct positive hit on the organoid well. **D)** Autofocus false positive from debris on the bottom of the PDMS membrane. **E)** Autofocus false positive from the top of the reservoir. **F)** Correct channel autofocus resulting from both sharp edges and Hough transform lines in the expected direction of the channel.

In addition to locating the circular central organoid well, the microscope had to autofocus on the bottom plane of the cells in the chip. When imaging the organoid well, this was complicated by the fact that the 2 mm deep circular well would generate a sharp image in any of those focal planes, that the top of the well connecting to the bottom surface of the reservoir could additionally be in focus, that the bottom surface of the PDMS membrane below the cells could generate a sharp image, that the cells themselves may be growing in 3D and generate sharp images in many focal planes. Additionally, the cells did not always generate particularly sharp images depending on how they were compacting and remodeling the extracellular matrix.

**Figure 9.**
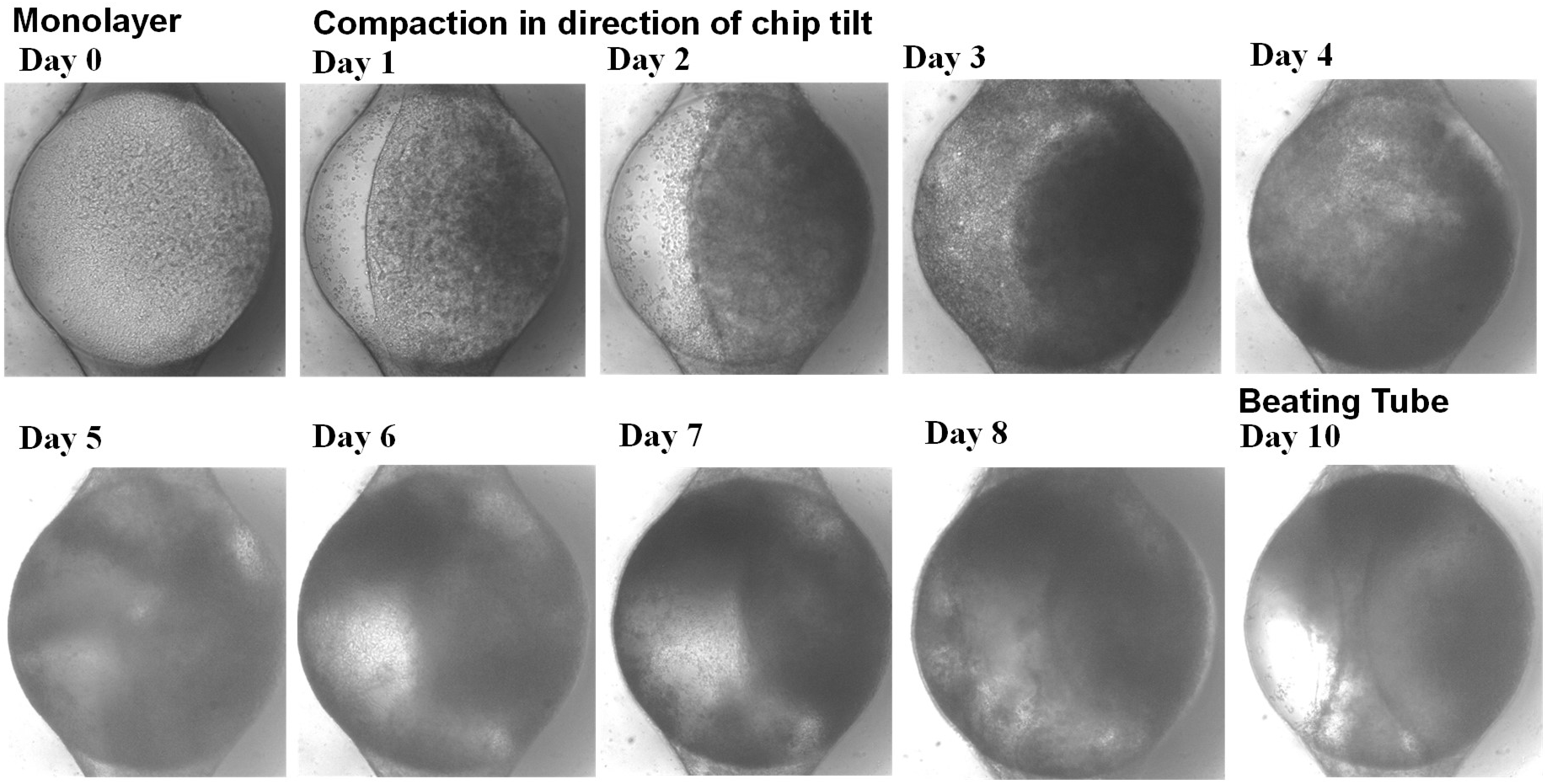
Time Lapse Imaging of an Organoid. A set of images were taken as iPSCs differentiated in situ from a pluripotent monolayer to a beating heart tube.

Autofocusing on the microfluidic channel, in contrast, was a simpler process with a defined structure that was only in focus within a narrow 100 μm range. This also made the channels harder to locate than the 2 mm high cylindrical well, which was visible in more focal planes. As such, the microscope was programmed to find the central organoid well first using Hough transforms, and then move by 1 mm to the location where the channel was expected to be in order to autofocus on the channel. The autofocus algorithm looked for both a sharp image and a good hit from a Hough line transform that looked for lines at the expected orientation of the channel, eliminating sharp images that might be focusing on the bottom surface of the PDMS membrane or the top surface of the reservoir (**Figure 7D-F**). This process had to be tolerant of angular error, as the users did not always position the chips precisely, and the chips sometimes shifted a little as the tilting stage moved. Angular errors in chip orientation were additionally calibrated as time lapse imaging was performed on each well, a necessary step as a small angular error could lead to a large position error between two distant wells.

Having focused on the microfluidic channel, the gantry then returned to the 800 μm organoid well, recentered on it, and began imaging the cells. Time lapse imaging included images of the cells in the organoid well and cells in the connecting channels.

An example set of time lapse images is shown in **Figure 8**. Images started the day after seeding, when the cells had formed a monolayer biased in the direction of the chip’s tilt. They then compacted in the direction of the bias, differentiated and proliferated, and eventually formed a beating heart tube.

## Discussion

As experiment after experiment was performed in order to understand and control differentiating heart organoids in a microfluidic environment, a brute force strategy of increasing the throughput of chip fabrication and differentiation experiments became necessary. This led to longer hours at microscopes, carrying chips back and forth to the incubator became a bottleneck for the project, and was additionally sub-optimal as it is preferable to study organoids at physiological temperatures and CO_2_ concentrations. It became apparent that a robotic system would be desirable for studying the organoids. A robotic microscope that could study the organoids as they developed, as well as pacing the organoids and controlling their tilt angle, became an integral part of the project.

Since there was no organoid imaging system that was affordable and satisfactory, a custom system was designed. After considering a wide variety of gantries, an architecture was selected that would be precise, stable, and could be constructed with off-the-shelf components or 3D printed parts. The microscope was assembled within a miniature aluminum extrusion frame to allow for easy modifications and alignment of optical components.

While the objective and tube lens selections were straightforward, the camera system was upgraded to a global shutter camera with a higher frame rate, necessary upgrades for studying beating cardiomyocytes and flow patterns. A fluorescent filter block was added that was based on a Semrock filter set that could implement three color fluorescent imaging without any moving parts. The system evolved over time, gaining a tilting stage that could generate gravitational biases and transient pressure heads, and an electrical pacing system that was flexible enough to handle a variety of simultaneous waveforms for both acute and chronic pacing.

As the system began collecting daily data on the organoid chips, it soon became clear that even a perfectly accurate gantry could not take images in repeatable locations if the chips were being moved by users and by the tilting stage. To accommodate this important real-world problem, an OpenCV machine vision system was added that was tailored to the microfluidic features and could find and image organoids despite misalignments by users.

